# ClOneHORT: Approaches for Improved Fidelity in Generative Models of Synthetic Genomes

**DOI:** 10.1101/2024.06.25.600651

**Authors:** Roland Laboulaye, Victor Borda, Shuo Chen, Kari E. North, Robert Kaplan, Timothy D. O’Connor

## Abstract

**Motivation:** Deep generative models have the potential to overcome difficulties in sharing individual-level genomic data by producing synthetic genomes that preserve the genomic associations specific to a cohort while not violating the privacy of any individual cohort member. However, there is significant room for improvement in the fidelity and usability of existing synthetic genome approaches.

**Results:** We demonstrate that when combined with plentiful data and with population-specific selection criteria, deep generative models can produce synthetic genomes and cohorts that closely model the original populations. Our methods improve fidelity in the site-frequency spectra and linkage disequilibrium decay and yield synthetic genomes that can be substituted in downstream local ancestry inference analysis, recreating results with .91 to .94 accuracy.

**Availability:** The model described in this paper is freely available at github.com/rlaboulaye/clonehort.

## 1 Introduction

As computational methods in genomics increasingly rely on biobank-level data, the need for large, accessible, and diverse genomic datasets increases [Byrska-Bishop et al., 2022, Taliun et al., 2021, All of Us Research Program Genomics Investigators, 2024, Sudlow et al., 2015, Sohail et al., 2023], which in turn presents significant challenges. For a multitude of reasons, genomic data cannot always be made publicly available. Even when it is made available, the products of large-scale initiatives to combine data are difficult to make widely accessible due to constraints in applications for the data and limited consent from participants of the original studies. At the same time, genomics experts need the inclusion of underrepresented populations to make a more comprehensive catalog of human variation, disease mapping, and personalized medicine for people everywhere. While some studies can be conducted on summary statistics that can be transferred without violating consent [Bodea et al., 2016, Borda et al., 2023, Artomov et al., 2024], some algorithms, such as association testing using mixed model approaches, local ancestry inference, and methods development, rely on individual-level genotype data.

To overcome these conflicting issues, recent research has proposed using neural networks trained on real genotype data, which often cannot be shared, to produce synthetic genomes that could be more widely distributed. This prior work has demonstrated that deep learning models are capable of generating synthetic genomes that mimic real data, in some cases well enough for limited use in downstream applications. Variational autoencoders (VAE) [Montserrat et al., 2019, Geleta et al., 2022], generative adversarial networks (GAN) [Montserrat et al., 2019, Yelmen et al., 2021, 2023], and generative moment matching networks (GMMN) [Perera et al., 2022] have shown varying degrees of success in producing synthetic genomes, often leveraging ancestry-conditioning to simplify disentangling latent representations and to enable ancestry-conditioned generation. In order to advance the state of synthetic genomes and build towards usability and adoption in real-world applications, we identify the following limitations:

1. Only non-admixed population modeling has been used to simplify labels of ancestry [Lewis et al., 2022] rather than also modeling populations with more complex demographic histories.
2. There is a lack of fidelity in the produced genotypes, which can be detected by the joint site frequency spectrum, introduced by the final step of the continuous-valued generative models intended to output discrete variant data, which in the case of the VAE-GAN [Montserrat et al., 2019] is a rounding operation. The effects are shown in Figure 1A.
3. There is a lack of fidelity in the produced genotypes at the individual level due to existing models failing to completely capture the local correlations of each individual’s genomic variants. These correlations contribute to the modeled population’s linkage disequilibrium (LD), and the modeling discrepancy can be expressed in terms of LD magnitude, as shown in Figure 1B.

**Figure 1.**
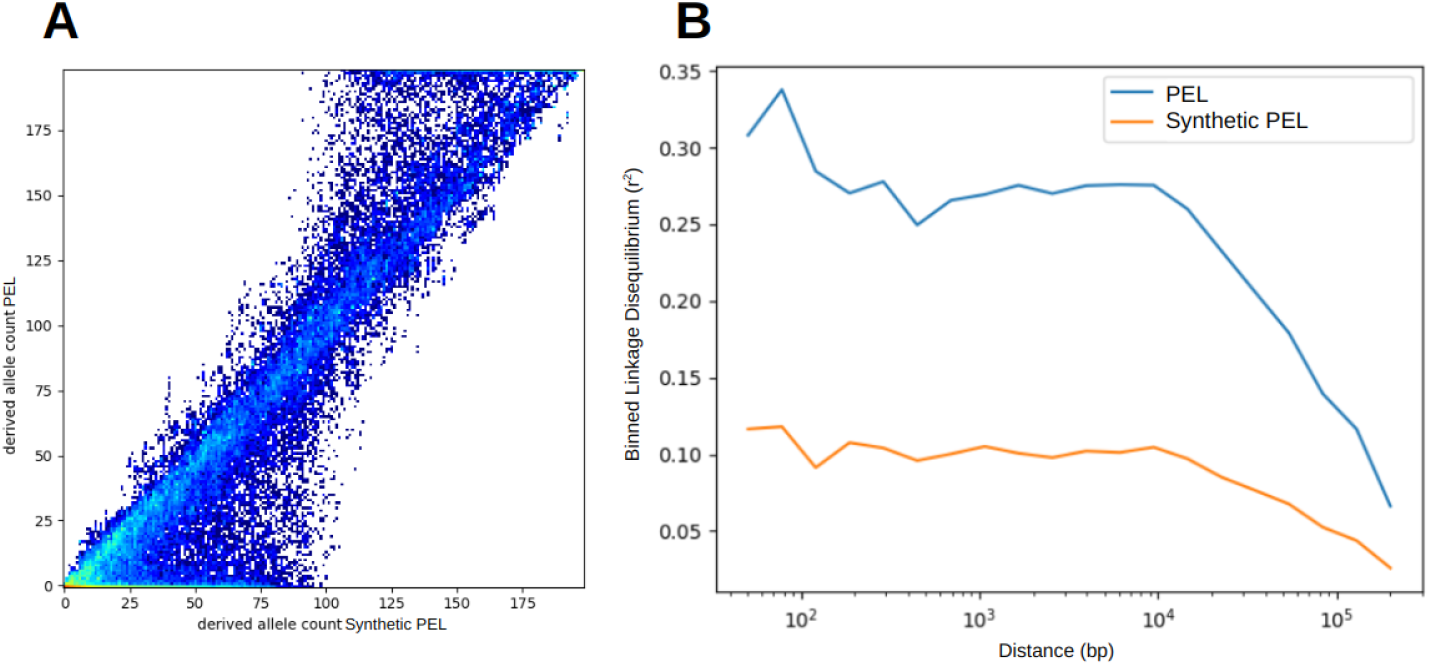
Prior work demonstrating capabilities and limitations of earlier models, in this case the model outlined in Montserrat et al. [2019] that we trained on 1000 Genomes Peruvians from Lima (PEL) [Consortium et al., 2015, Byrska-Bishop et al., 2022]. A) Joint site frequency spectrum between real and synthetic Peruvian 1000 Genomes population, showing signification deviation, and B) Binned LD difference between real and synthetic Peruvian 1000 Genomes population.

Building on prior efforts and recognizing that widespread adoption of synthetic genomes will require overcoming well-founded skepticism, we introduce a set of approaches aimed at modeling a wider variety of populations and improving the fidelity of produced synthetic genomes to the original samples according to key genomic metrics, while preserving the anonymity of individual genomic data of real people. We present the following contributions:

1. We leverage admixed individuals to increase the size of our training pool and to allow our model to better capture the spectrum of diversity represented by each population we try to model, taking as our focus Latin American individuals with predominantly three-way admixture [Manichaikul et al., 2012, Bryc et al., 2010].
2. We alter the generative model’s training criteria, by using a Negative-Binomial Variational Autoencoder (NBVAE) [Zhao et al., 2020] to better account for genotype data’s discrete nature.
3. We develop a simulated-annealing-based selection procedure where we can oversample synthetic genomes from our generative model and isolate the subset of genomes that most faithfully represents key evolutionary statistics for the population we are modeling.

## 2 Materials and Methods

We begin by outlining our evaluation methodology and proceed to detail our expanded data pool, our model architecture, and our selection process.

### 2.1 Deepening Evaluation Methodology

Given the research context in which synthetic genomes have the potential to be used, it is incumbent upon us to perform an extensive and multifaceted evaluation of their quality. We want to generate synthetic genomes that look real, can be used in place of real genomes, and don’t violate the privacy of the genomes on which the generative model was trained. We outline below the various evaluation metrics we will use to encompass the aforementioned goals:

1. Look real: Since this is a broad goal, we will use a variety of metrics to compare the synthetic genomes to the real genomes at the population statistic level. We will do so by comparing the site frequency spectrum, linkage disequilibrium decay, and PCA-projected visualizations of the genomes.
2. Be used like real genomes: The purpose of synthetic genomes is to substitute real genomes in applications that require individual-level genomic data. For this evaluation, we will use Local Ancestry Inference (LAI) as our test application. We begin by choosing the reference population that we want to model. Then, using that population as one of *k* reference populations, we generate LAI calls for an admixed population as our truth set. We then repeat that process, only this time substituting the synthetic version of the population of interest as one of our reference populations. The LAI calls on the admixed population generated by this second process are compared to the truth set and the comparison is reported as local ancestry replication.
3. Don’t violate privacy: Even if generated synthetic genomes perform well on the metrics outlined in the prior two sections, they cannot be shared if they violate the privacy of the individuals whose genomic data was used to train the generative model. In order to evaluate privacy protection, we will use a privacy loss approach, shown in Equation 1, adapted from prior work proposed by Yale [Dash et al., 2019] and applied to synthetic genomes by Yelmen [Yelmen et al., 2021, 2023].

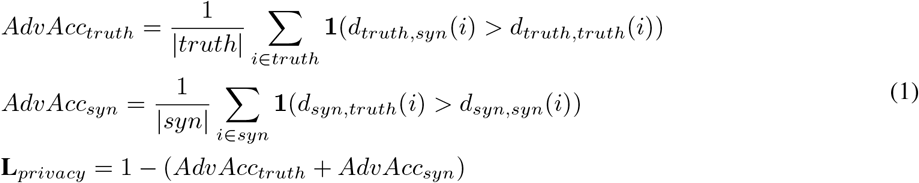

The adversarial accuracy values defined in Equation 1 measure, according to a nearest neighbor distance metric *d* defined over genomes (such as genomic distance or Mahalanobis distance between PCA-projections), two aspects of similarity: 1) the proportion of the real genomes’ nearest neighbors that lie in the real rather than synthetic data, and 2) the proportion of the synthetic genomes’ nearest neighbors that lie in the synthetic rather than real data. Using *d*, we define *d*_*truth*,*syn*_(*i*) and *d*_*truth*,*truth*_(*i*) to be functions representing the distance from a genome in the truth set (a real genome) and its nearest neighbor in the synthetic set and the truth set respectively. *d*_*syn*,*truth*_(*i*) and *d*_*syn*,*syn*_(*i*) are similar, but are defined over genomes in the synthetic set. When *d*_*truth*,*syn*_(*i*) *> d*_*truth*,*truth*_(*i*), it means the *i*^*th*^ real genome more closely resembles another real genome than a synthetic genome. If true for the majority of the dataset, this metric suggests that the model may not be adequately capturing the real genomes’ distribution. If *d*_*truth*,*syn*_(*i*) *< d*_*truth*,*truth*_(*i*) for a majority of real genomes, this may suggest that the model is too closely copying each individual real genome, in which case sharing the synthetic genomes may violate the privacy of those to whom the real genomes belong. Achieving an *AdvAcc*_*truth*_ value of near 0.5 would indicate that the model is balancing between these two extremes, producing convincing synthetic genomes that preserve privacy.

In the case of *d*_*syn*,*truth*_(*i*) and *d*_*syn*,*syn*_(*i*), *d*_*syn*,*truth*_(*i*) *< d*_*syn*,*syn*_(*i*) for the majority of the synthetic genomes also suggests that privacy may be violated, while *d*_*syn*,*truth*_(*i*) *> d*_*syn*,*syn*_(*i*) might indicate that the synthetic genomes have bunched together in clumps, possibly indicating that the model is suffering from mode collapse. Therefore, the optimal value for *AdvAcc*_*syn*_ is also 0.5. The adversarial accuracies for real and synthetic genomes are combined in **L**_*privacy*_, the privacy loss. Note that while 0.5 is the optimal value for both adversarial accuracies, since the privacy loss only cares about adversarial accuracy values that imply copying, it only penalizes low values of adversarial accuracy. Since high values are indicative of low fidelity, our other metrics will suffice.

### 2.2 Enlarged Data Pool

Prior work has focused entirely on modeling non-admixed populations [Montserrat et al., 2019, Geleta et al., 2022, Yelmen et al., 2023]. This is limiting in two ways. First, a large portion of available training data from large consortiums such as the Trans-Omics for Precision Medicine (TOPMed) project [Taliun et al., 2021], data on which deep learning models are highly dependent, contains admixed individuals. Second, the genomic datasets for which synthetic genome approaches would prove useful, those that could be used in downstream applications but cannot be disseminated for privacy concerns, will often be admixed. As such, synthetic genome approaches should be built to leverage and model both admixed and non-admixed populations.

Our generative model is class conditional, which means that it accepts continental ancestry population labels to condition the probability distribution and sample synthetic genomes. The model accepts these labels for each window along the genome. This allows us to leverage non-admixed populations, for which every window will have the same label, as well as admixed populations, for which continental ancestry labels will vary along the genome. We leverage reference populations and Latin American populations from 1000 Genomes [Consortium et al., 2015, Byrska-Bishop et al., 2022], comprising 1570 individuals; and the Peruvian Genome Project (PGP) [Harris et al., 2018], comprising 148 individuals; and also augment our training set with 8,755 Latin American individuals from the Hispanic Community Health Study / Study of Latinos (HCHS/SOL) Project [Feofanova et al., 2020] obtained in collaboration through TOPMed [Taliun et al., 2021]. Unlike the continental ancestry reference populations we used, the Latin American genomes, as well as other admixed individuals, underwent local ancestry inference as preprocessing to generate continental ancestry labels for every genotype window. This local ancestry inference was performed using the Gnomix software [Hilmarsson et al., 2021] and with Utahn individuals (CEU 1000 Genomes; N=184), Yoruba individuals from Ibadan (YRI 1000 Genomes; N=120), and individuals with a high proportion of Native American ancestry from PGP (N=148) [Harris et al., 2018] as continental reference populations. We expand on prior work by accepting conditioning labels, i.e. the posterior probabilities from the local ancestry inference, for each window, allowing us to train our model on admixed genomes and accommodate the potential error.

### 2.3 Model Architecture

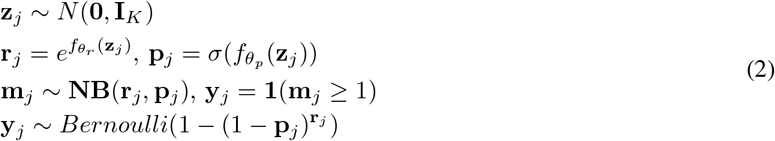

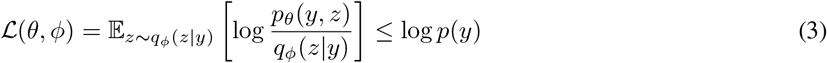

We build on the approach used in Mas Montserrat et al. [Montserrat et al., 2019], which used a windowed variational autoencoder coupled with a generative adversarial network (VAE-GAN) [Larsen et al., 2016]. While that model consists of three network components, an encoder, decoder, and discriminator, we simplify the modeling procedure by removing the discriminator, which, as suggested by Limitation 2, offered few benefits since the post-decoder rounding operation does not allow for backpropagation, preventing the discriminator from operating on the real output data. Instead, following our decoder, we opt to apply a Bernoulli sampling procedure, which like rounding introduces its own lack of fidelity that we address by replacing the VAE [Kingma and Welling, 2013] with a Negative-Binomial VAE (NBVAE) [Zhao et al., 2020].

Equation 2 describes the generative process that underpins our model. As is typical for VAE, we assume a normally-distributed, *K*-dimentional latent space with mean 0 and an identity covariance matrix *I*_*K*_, from which we draw a latent vector ***z***_*j*_. The decoder, parameterized by ***θ*** (***θ***_*r*_ and ***θ***_*p*_) and represented by 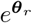 and *σ*(***θ***_*p*_), maps ***z***_*j*_ to two outputs, ***r***_*j*_ and ***p***_*j*_ . These outputs are in turn used to parameterize a Negative Binomial distribution (NB) from which ***m***_*j*_ is drawn, where ***m***_*j*_ is the parameter of a Bernoulli distribution over the discrete value of interest ***y***_*j*_ , a binary value representing the presence or absence of a variant at location *j*. Equation 2’s final line shows that ***m***_*j*_ can be marginalized out and the NBVAE’s output ***y***_*j*_ can be represented as a Bernoulli probability that depends only on rj and pj. Compared to a typical VAE, this formulation gives the model finer control over the output probability than a sigmoid function, producing more exact probabilities and reducing the error introduced by post-decoder sampling.

The NBVAE is trained by minimizing the Evidence-Lower-Bound Loss shown in Equation 3, where *z* is the latent vector, *y* is the variant data, and ***θ*** and ***ϕ*** represent parameters of the decoder and encoder respectively. Both the encoder and decoder are composed of multi-layer perceptrons, containing 3 hidden layers, each with 512 nodes. We train a separate NBVAE model for each 0.2*cM* window on each chromosome. We outline our model layout in Figure 2A.

**Figure 2.**
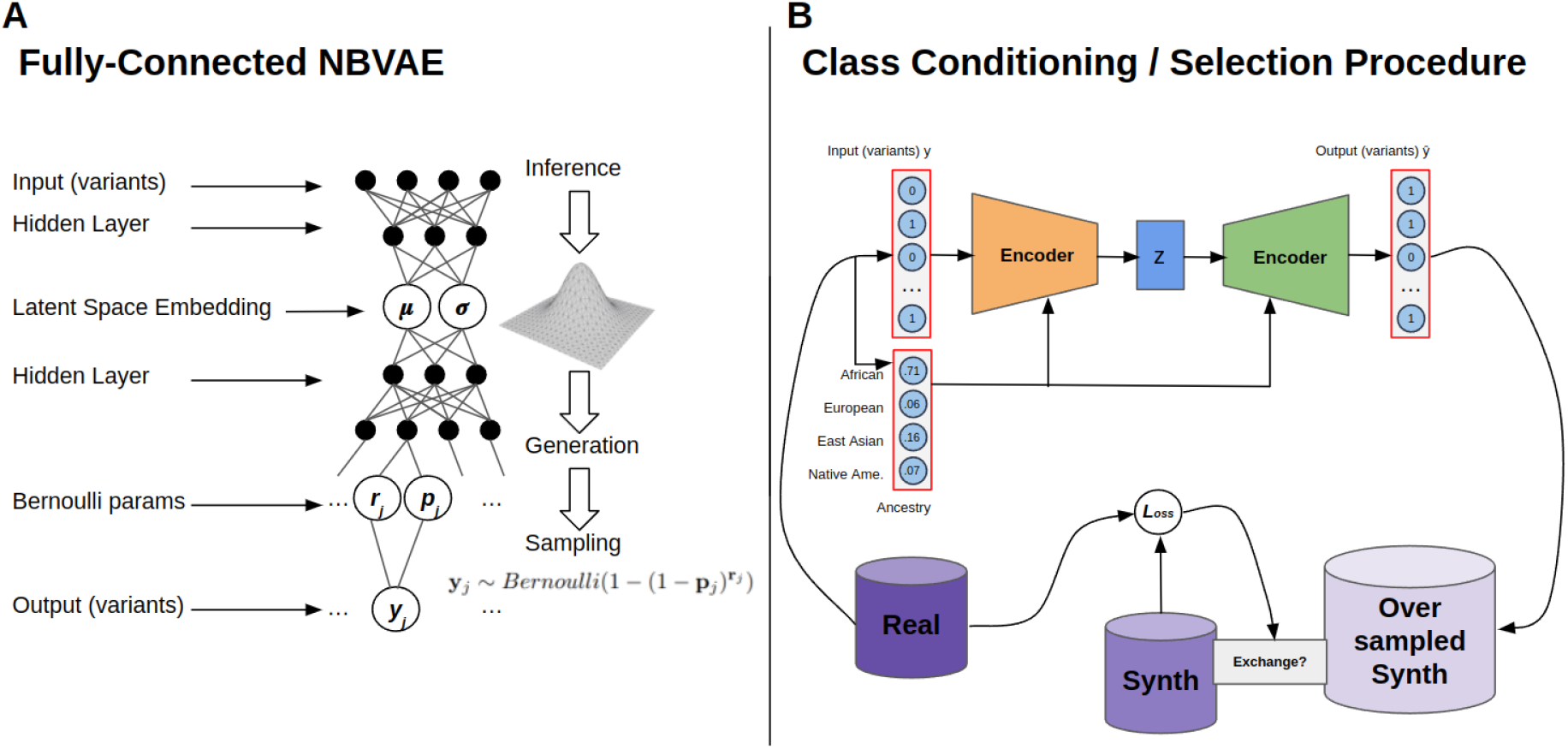
Generative model layout. A) The diagram shows the general encoder-decoder layout of the NBVAE. Note that the latent space in which the genomes are embedded is assumed to be Gaussian. B) The diagram gives a rough sketch of the class-conditioning process by which the NBVAE is trained to associate continental labels with specific genotype patterns. It also illustrates the simulated annealing process by which synthetic genomes are selected from the oversampled set based on the way they compare to real genomes.

### 2.4 Selection Procedure

In order to further address Limitations 2 and 3, we develop a selection procedure whereby we can oversample synthetic genomes from our generative model and isolate the genomes that best fit our evaluation criteria. To that end, we use the loss function outlined in Equation 4. The first three lines of the equation define the three components of the loss, designed to ensure that the selected genomes match the real genomes in genotype distribution and linkage disequilibrium while preserving privacy. ***L***_*genotype*_ compares the allele frequencies of the synthetic cohort with those of the real cohort, where *n* corresponds to the number of SNPs. ***L***_*linkage*_ is defined as the average of the difference in *r*^2^ (a linkage disequilibrium metric) between the real and synthetic genomes is highest. However, since calculating linkage disequilibrium for every iteration of our selection procedure would be too computationally costly, rather than using all SNP pairs within windows, we select a subset of pairs where the difference in *r*^2^ is highest. ***L***_*privacy*_ implements a variation on the privacy loss described in greater detail in Equation 1. We can’t directly use the privacy loss as previously described for this local optimization because it doesn’t account for replication (but we use privacy loss from Equation 1 in our later evaluation). These three loss components can be combined with different coefficients *α, β*, and *γ* based on which the user most wants to emphasize.

We optimize for the loss ***L*** using simulated annealing [Kirkpatrick et al., 1983]. The selection procedure consists of oversampling by some factor, randomly selecting a starting set of synthetic genomes, and iteratively selecting a neighboring set, evaluating its loss according to Equation 4, using both the loss and iteration temperature to determine whether to accept the new set. We define a neighboring set as one with *m* replaced synthetic individuals (we used *m* = 20).

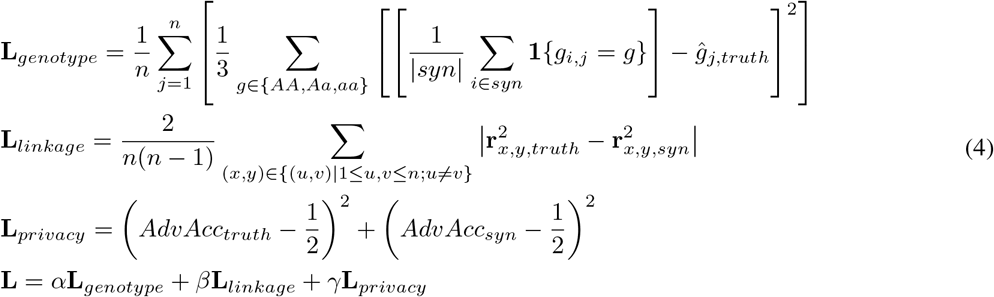

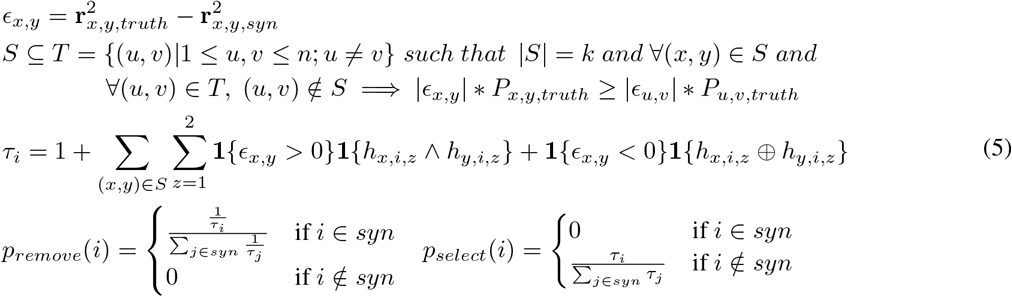

The neighborhood sampling function is outlined in Equation 5 and it prioritizes adding to the set of genomes that contribute positively to the LD modeling. This simulated annealing selection procedure ensures that our synthetic genomes, while diverse, more closely match the real genomes’ allele frequency and LD patterns. A simple illustration is designated in Figure 2B as “Exchange?”.

## 3 Results

The following results were obtained by training the NBVAE with and without augmentation and the selection procedure, on chromosomes 20, 21, and 22 on the data described in Section 2.2. As outlined in Section 2.1, we first focus our evaluation on measuring how well the generated genomes capture the population of interest’s distribution. In Figure 3, the joint site frequency spectrum demonstrates that the synthetic genomes match the allele frequencies of the original population quite closely. The leftmost column presents joint site frequencies with three real populations from 1000 Genomes, Esan individuals from Nigeria (ESN), British individuals from England and Scotland (GBR), and Peruvians from Lima, Peru (PEL) to compare with YRI, CEU, and PGP respectively. As evidenced by the narrowing in the joint site frequency spectrum from the left plots to the right, our produced synthetic genomes match allele frequencies better than other real populations that share continental ancestry. Adding the selection procedure further tightens the joint site frequency spectrum, indicating more faithful replication of the original population’s allele frequencies. Figure 4 demonstrates that the magnitude and spread of linkage disequilibrium of a synthetic version of PGP matches that of the real PGP, significantly closing the gap observed in prior work (see Figure 1). The selection procedure enables a large jump in fidelity, allowing the model to produce synthetic genomes that more closely match LD than a similar real population, in this case PEL.

**Figure 3.**
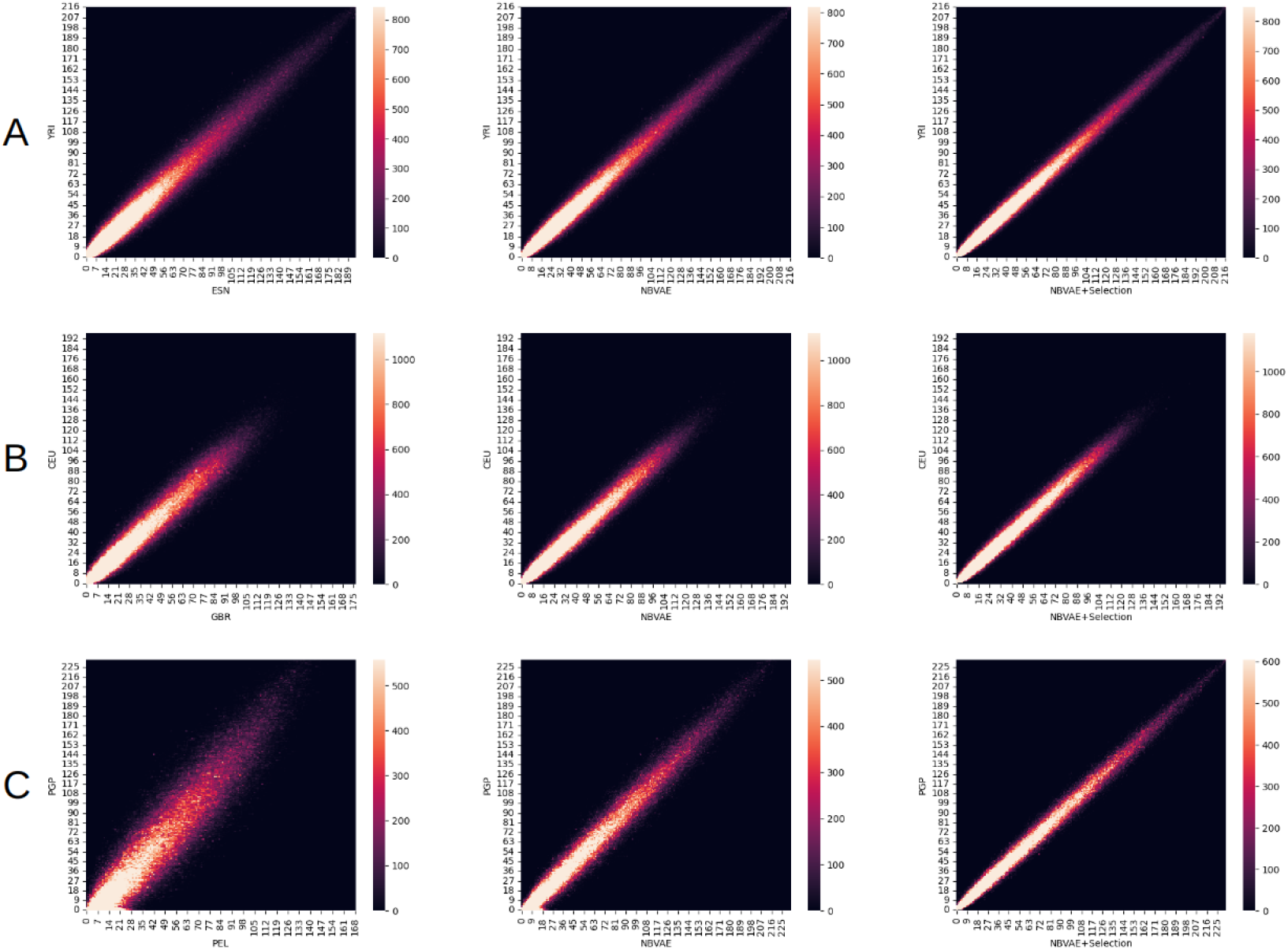
Joint site frequency spectra demonstrating our model’s ability to capture genetic variation, using a joint site frequency spectrum with a similar real cohort as a baseline. **A** compares the Yoruba cohort from Nigeria to the Esan Cohort from Nigeria, both from the 1000 Genomes Project, and to synthetic versions of the Yoruba cohort generated with an NBVAE and an NBVAE followed by a selection procedure respectively. **B** presents the same using Utahns of European descent and Britains, both from 1000 Genomes, as reference and similar real population. **C** presents the same using individuals with predominantly Native American ancestry from the Peruvian Genome Project as the reference and Peruvians from Lima, also from 1000 Genomes, as the similar real population.

**Figure 4.**
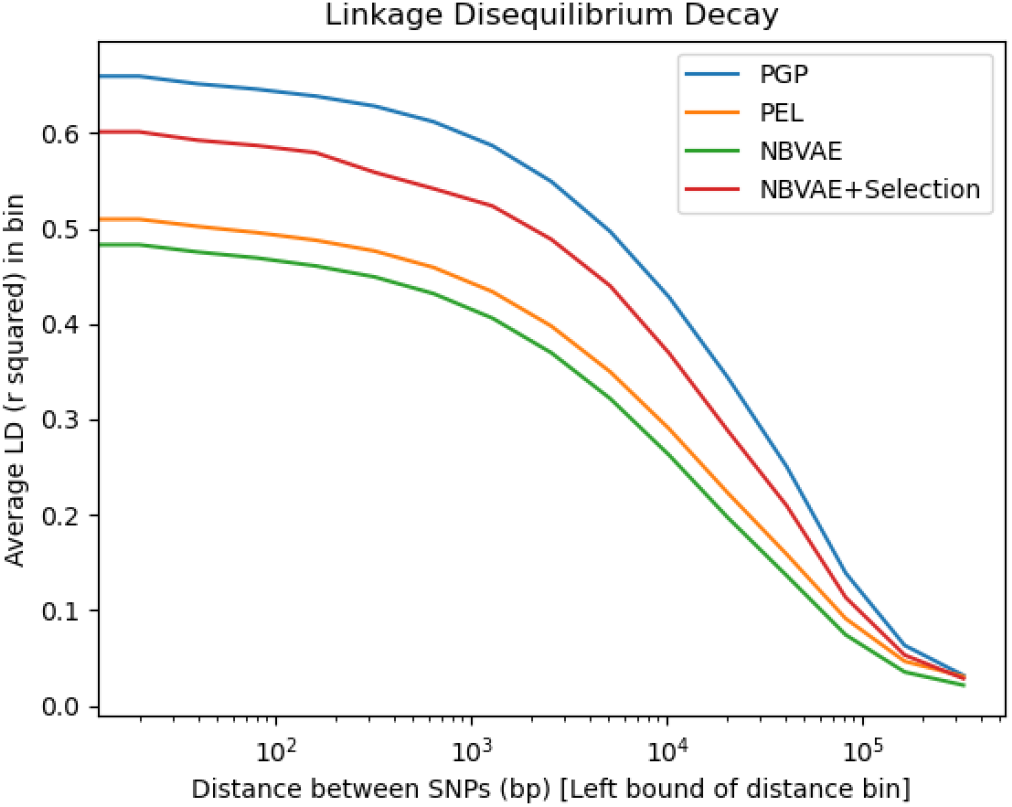
Improvement in linkage disequilibrium modeling when using selection procedure after NBVAE. The models are trained on PGP and PEL is included for comparison.

A visual inspection of Figure 5 further demonstrates that the synthetic genomes, when projected onto a shared PCA space with the original samples, capture every portion of the population of interest. The fairly evenly distributed density suggests that the generative models are outputting a diverse set of synthetic individuals rather than a set of repetitive individuals. For this evaluation, the selection procedure appears to have limited effect, except perhaps to slightly tighten the distribution. Note that PEL in Figure 5C fails to emulate the distribution of PGP, but this is to be expected since PEL is comprised of admixed individuals of Indigenous American (average of 60%), European, and African ancestries living in Lima, unlike PGP which is comprised of individuals of primarily Indigenous American ancestry (average of 90%) from regions across North and South Peru. The discrepancy between PGP and PEL highlights the uniqueness of PGP as a resource and the resulting benefit that would come from being able to access a synthetic version of that cohort.

**Figure 5.**
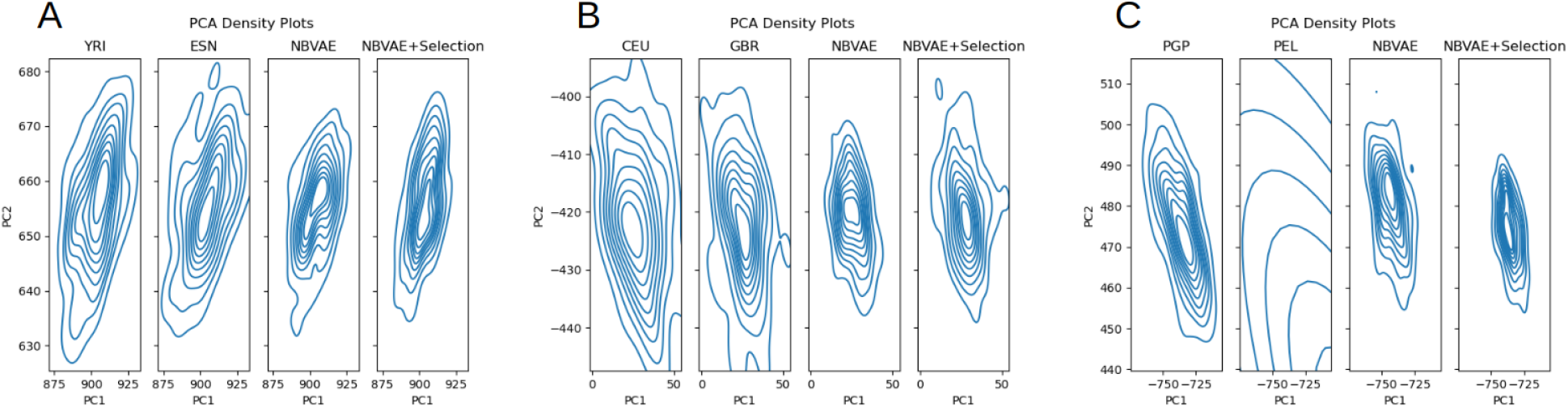
PCA density plots showing how closely our synthetic genomes mirror the target populations’ distribution in PCA space, as compared to similar real populations. **A** compares the Yoruba cohort from Nigeria to the Esan Cohort from Nigeria, both from the 1000 Genomes Project, and to synthetic versions of the Yoruba cohort generated with an NBVAE and an NBVAE followed by a selection procedure respectively. **B** presents the same using Utahns of European descent and Britains, both from 1000 Genomes, as reference and similar real population. **C** presents the same using individuals with predominantly Native American ancestry from the Peruvian Genome Project as the reference and Peruvians from Lima, also from 1000 Genomes, as the similar real population.

Following the pattern set in [Montserrat et al., 2019], we used local ancestry inference as a means of evaluating the synthetic genomes’ value in a downstream analysis. For the samples to be labeled, we used three admixed Latin American populations from the 1000 Genomes Project: Mexican Ancestry in Los Angeles (MXL), Colombian in Medellin (CLM), and Puerto Rican in Puerto Rico (PUR). For our reference populations, we used the aforementioned CEU and YRI populations and sequenced individuals with a high proportion of Native American ancestry from the Peruvian Genome Project (PGP) [Harris et al., 2018]. Since there are no true labels of continental ancestry for our admixed samples, we instead used the labels inferred when using all three real reference populations. We then replaced CEU with synthetic CEU, inferred continental ancestry, and compared the new labels with those generated using the real CEU, dubbing the comparison local ancestry replication accuracy. We repeated the experiment with real and synthetic PGP. The local ancestry inference was performed using Gnomix [Hilmarsson et al., 2021]. We report results from comparing windows where Gnomix produced a posterior probability for the label *>* 0.9 and also report results from comparing all windows in parentheses.

Tables 1 and 2 demonstrate how closely our synthetic genomes can replicate results one would get from using real genomes are a reference. The last row of each table shows results from using a comparable real population, GBR and PEL for CEU and PGP respectively, and highlights that substituting in a synthetic population as a reference population would alter local ancestry inference results by about the same amount that using an alternate real population of the same continental ancestry.

**Table 1:** Comparison of local ancestry replication values for synthetic genomes, intended to mimic the CEU 1000 genomes population, generated by different models and configurations. We compare their ability to, when used as a reference population replacing CEU, infer continental ancestry labels for 3 admixed Latin American populations from 1000 Genomes. GBR, a real 1000 Genomes population of British individuals from England and Scotland, is included as a real population baseline. The primary results reported compared only windows where the LAI produced a posterior probability of *>* 0.9 while the results in parentheses are for all windows.

| LAI Replication<br>Synthetic CEU |  |  | Population on which<br>LAI is Performed |  |  |
| --- | --- | --- | --- | --- | --- |
| Model | Augmentation | Selection | MXL | CLM | PUR |
| VAE-GAN | - | - | 0.860 (0.798) | 0.814 (0.765) | 0.827 (0.792) |
| NBVAE | × | × | 0.904 (0.861) | 0.911 (0.778) | 0.898 (0.800) |
| NBVAE | × | ✓ | 0.914 (.853) | 0.906 (0.791) | 0.901 (0.812) |
| NBVAE | ✓ | × | 0.902 (0.862) | 0.905 (0.784) | 0.900 (0.803) |
| NBVAE | ✓ | ✓ | **0.942** (0.881) | **0.924** (0.807) | **0.917** (0.843) |
| GBR | - | - | 0.949 (0.850) | 0.911 (0.812) | 0.918 (0.812) |

**Table 2:** Comparison of local ancestry replication values for synthetic genomes, intended to mimic the PGP population, generated by different models and configurations. We compare their ability to, when used as a reference population replacing PGP, infer continental ancestry labels for 3 admixed Latin American populations from 1000 Genomes. The primary results reported compared only windows where the LAI produced a posterior probability of *>* 0.9 while the results in parentheses are for all windows.

| LAI Replication<br>Synthetic PGP |  |  | Population on which<br>LAI is Performed |  |  |
| --- | --- | --- | --- | --- | --- |
| Model | Augmentation | Selection | MXL | CLM | PUR |
| VAE-GAN | - | - | 0.831 (0.802) | 0.809 (0.808) | 0.815 (0.817) |
| NBVAE | × | × | 0.887 (0.831) | 0.871 (0.852) | 0.857 (0.859) |
| NBVAE | × | ✓ | 0.905 (0.878) | 0.903 (0.880) | 0.898 (0.869) |
| NBVAE | ✓ | × | 0.892 (0.830) | 0.881 (0.847) | 0.854 (0.849) |
| NBVAE | ✓ | ✓ | **0.921** (0.900) | **0.915** (0.891) | **0.914** (0.886) |
| PEL | - | - | 0.918 (0.883) | 0.915 (0.882) | 0.909 (0.878) |

## 4 Discussion

As outlined in our introduction, the main challenges our methods seek to address are modeling admixed populations and improving modeling fidelity as demonstrated through joint site frequency spectra and linkage disequilibrium decay. The visual inspections provided in Figures 3, 4, 5 demonstrate how closely the synthetic genomes can recreate complex admixed populations. The inclusion of admixed, local-ancestry labeled training data ensures that all variation in the populations is captured, as demonstrated by the fact that our PCA plots do not include any areas lacking overlap. The improved NBVAE training criteria combined with the guided post-generation filtering process ensure that every synthetic genome is of high quality and that complex LD disequilibrium patterns in the original data are captured, significantly outperforming prior approaches. The selection criteria provides the vast majority of the improvement in LD decay modeling, suggesting that the pseudo-LD correction in Equation 4 is having the intended effect.

These benefits yielded further improvements when the genomes were used downstream for local ancestry replication for both European and Native American ancestry genomes. While each component of our approach combined to improve fidelity in the aforementioned comparisons, this was not true for every component when looking at LAI. For CEU LAI replication the NBVAE model provided an average .071 increase in accuracy over the prior VAE-GAN. The augmentation failed to notably affect results, but the selection criteria further increased the gain over the baseline to .094. For PGP LAI replication, the improvements are .053 and .098 respectively, once again demonstrating the importance of using not only realistic synthetic individuals but an entire realistic synthetic cohort.

Furthermore, the aforementioned improvements are achieved largely without compromising individual genome privacy, as demonstrated in Table 3. As mentioned in 2.1, ideal adversarial accuracy values for both *AdvAcc*_*truth*_ and *AdvAcc*_*syn*_ should be 0.5. For privacy loss, lower values indicate a lower chance of violating privacy. As demonstrated by its high values of *AdvAcc*_*truth*_, the VAE-GAN model produces genomes with slightly lower individual fidelity, which in turn lowers its privacy loss. The switch to NBVAE improves fidelity, but at the cost of increased *AdvAcc*_*syn*_, indicating some clumping around easier to model areas of the distribution. Adding in the selection procedure mostly preserves fidelity while diversifying the set of synthetic genomes, lowering *AdvAcc*_*syn*_. In all cases, the *AdvAcc* values stay above 0.5 and consequently privacy loss stays below 0, indicating that the models are not copying the real genomes. Thus, NBVAE+Selection’s improvements detailed in prior sections do not come at the cost of violating privacy.

**Table 3:**
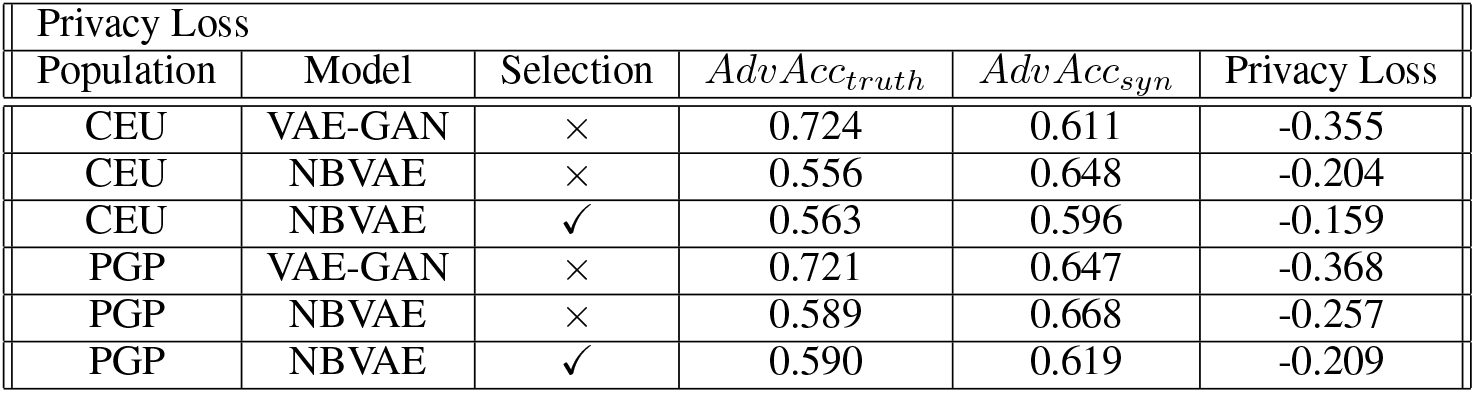
Comparison of privacy loss (Equation 1) values for synthetic genomes generated by different models and configurations. We compare the privacy loss for synthetic CEU and PGP populations generated with a VAE-GAN, the NBVAE model, and the NBVAE model passed through a selection procedure.

| Privacy Loss |  |  |  |  |  |
| --- | --- | --- | --- | --- | --- |
| Population | Model | Selection | $AdvAcc_{truth}$ | $AdvAcc_{syn}$ | Privacy Loss |
| CEU | VAE-GAN | × | 0.724 | 0.611 | -0.355 |
| CEU | NBVAE | × | 0.556 | 0.648 | -0.204 |
| CEU | NBVAE | ✓ | 0.563 | 0.596 | -0.159 |
| PGP | VAE-GAN | × | 0.721 | 0.647 | -0.368 |
| PGP | NBVAE | × | 0.589 | 0.668 | -0.257 |
| PGP | NBVAE | ✓ | 0.590 | 0.619 | -0.209 |

However, our method’s ability to model genomic data works only for somewhat common variants, and attempts to model data with minor allele frequency greater than 0.001 as opposed to 0.01 performed poorly, demonstrating that our current generation pipeline has a limit when it comes to modeling complex LD patterns between rare variants. This can be concerning since the sort of small populations that would be beneficial to synthesize and share are in part of interest because of their unique rare variants. We address potential approaches to remedy this issue in the next section.

## 5 Conclusion

### 5.1 Summary

We presented a set of approaches meant to improve the quality of synthetic genomes generated by deep learning models. We demonstrated that all three approaches contribute to improvements in individual genome quality, cohort-level quality, and downstream performance. Most notably, we demonstrated that guided post-generation filtering (our selection procedure) is an essential step in making synthetic genomes usable.

### 5.2 Limitations and Future Work

As shown in our results, our current generation pipeline generates significantly less faithful genomes when the minor allele frequency threshold used to filter the data of interest is lowered. We expect that larger amounts of training data will be required to properly model the larger sequences that lower minor allele frequency thresholds yield. Furthermore, we suspect that adopting a neural network architecture capable of jointly modeling larger sections of the variant sequences at a time would also yield increased fidelity. To that end, we will try to incorporate large convolutional layers and / or patches under a transformer in our NBVAE.

## Acknowledgements

We gratefully acknowledge the studies and participants who provided biological samples and data for the Hispanic Community Health Study / Study of Latinos and the Peruvian Genome Project. We also acknowledge the Trans-Omics for Precision Medicine (TOPMed) for enabling this collaboration.

## References

Marta Byrska-Bishop, Uday S Evani, Xuefang Zhao, Anna O Basile, Haley J Abel, Allison A Regier, André Corvelo Wayne E Clarke, Rajeeva Musunuri, Kshithija Nagulapalli, et al. High-coverage whole-genome sequencing of the expanded 1000 genomes project cohort including 602 trios. Cell, 185(18):3426–3440, 2022.

Daniel Taliun, Daniel N Harris, Michael D Kessler, Jedidiah Carlson, Zachary A Szpiech, Raul Torres, Sarah A Gagliano Taliun, André Corvelo Stephanie M Gogarten, Hyun Min Kang, et al. Sequencing of 53,831 diverse genomes from the nhlbi topmed program. Nature, 590(7845):290–299, 2021.

placeholder All of Us Research Program Genomics Investigators. Genomic data in the all of us research program. Nature, 627(8003):340, 2024.

Cathie Sudlow, John Gallacher, Naomi Allen, Valerie Beral, Paul Burton, John Danesh, Paul Downey, Paul Elliott, Jane Green, Martin Landray, et al. Uk biobank: an open access resource for identifying the causes of a wide range of complex diseases of middle and old age. PLoS medicine, 12(3):e1001779, 2015.

Mashaal Sohail, María J Palma-Martínez, Amanda Y Chong, Consuelo D Quinto-Cortés, Carmina Barberena-Jonas, Santiago G Medina-Muñoz, Aaron Ragsdale, Guadalupe Delgado-Sánchez, Luis Pablo Cruz-Hervert, Leticia Ferreyra-Reyes, et al. Mexican biobank advances population and medical genomics of diverse ancestries. Nature, 622(7984):775–783, 2023.

Corneliu A Bodea, Benjamin M Neale, Stephan Ripke, Murray Barclay, Laurent Peyrin-Biroulet, Mathias Chamaillard, Jean-Frederick Colombel, Mario Cottone, Anthony Croft, Renata D’Incà, et al. A method to exploit the structure of genetic ancestry space to enhance case-control studies. The American Journal of Human Genetics, 98 (5):857–868, 2016.

Victor Borda, Douglas P Loesch, Bing Guo, Roland Laboulaye, Diego Veliz-Otani, Jennifer N French-Kwawu, Thiago Peixoto Leal, Stephanie M Gogarten, Sunday Ikpe, Mateus H Gouveia, et al. Genetics of latin american diversity (glad) project: insights into population genetics and association studies in recently admixed groups in the americas. bioRxiv, pages 2023–01, 2023.

Mykyta Artomov, Alexander A Loboda, Maxim N Artyomov, and Mark J Daly. Public platform with 39,472 exome control samples enables association studies without genotype sharing. Nature Genetics, pages 1–9, 2024.

Daniel Mas Montserrat, Carlos Bustamante, and Alexander Ioannidis. Class-conditional vae-gan for local-ancestry simulation. arXiv preprint 1911.13220, 2019.

Margarita Geleta, Daniel Mas Montserrat, Carlos Bustamante, X Giró-i Nieto, and Alexander Ioannidis. Deep variational autoencoders for population genetics. biorxiv, 2022.

Burak Yelmen, Aurélien Decelle, Linda Ongaro, Davide Marnetto, Corentin Tallec, Francesco Montinaro, Cyril Furtlehner, Luca Pagani, and Flora Jay. Creating artificial human genomes using generative neural networks. PLoS genetics, 17(2):e1009303, 2021.

Burak Yelmen, Aurélien Decelle, Leila Lea Boulos, Antoine Szatkownik, Cyril Furtlehner, Guillaume Charpiat, and Flora Jay. Deep convolutional and conditional neural networks for large-scale genomic data generation. PLOS Computational Biology, 19(10):e1011584, 2023.

Maria Perera, Daniel Mas Montserrat, Miriam Barrabes, Margarita Geleta, Xavier Giró-i Nieto, and Alexander G Ioannidis. Generative moment matching networks for genotype simulation. In 2022 44th Annual International Conference of the IEEE Engineering in Medicine & Biology Society (EMBC), pages 1379–1383. IEEE, 2022.

Anna CF Lewis, Santiago J Molina, Paul S Appelbaum, Bege Dauda, Anna Di Rienzo, Agustin Fuentes, Stephanie M Fullerton, Nanibaa’A Garrison, Nayanika Ghosh, Evelynn M Hammonds, et al. Getting genetic ancestry right for science and society. Science, 376(6590):250–252, 2022.

Ani Manichaikul, Walter Palmas, Carlos J Rodriguez, Carmen A Peralta, Jasmin Divers, Xiuqing Guo, Wei-Min Chen, Quenna Wong, Kayleen Williams, Kathleen F Kerr, et al. Population structure of hispanics in the united states: the multi-ethnic study of atherosclerosis. PLoS genetics, 8(4):e1002640, 2012.

Katarzyna Bryc, Christopher Velez, Tatiana Karafet, Andres Moreno-Estrada, Andy Reynolds, Adam Auton, Michael Hammer, Carlos D Bustamante, and Harry Ostrer. Genome-wide patterns of population structure and admixture among hispanic/latino populations. Proceedings of the National Academy of Sciences, 107(supplement_2): 8954–8961, 2010.

He Zhao, Piyush Rai, Lan Du, Wray Buntine, Dinh Phung, and Mingyuan Zhou. Variational autoencoders for sparse and overdispersed discrete data. In International Conference on Artificial Intelligence and Statistics, pages 1684–1694. PMLR, 2020.

1000 Genomes Project Consortium et al. A global reference for human genetic variation. Nature, 526(7571):68, 2015.

Saloni Dash, Ritik Dutta, Isabelle Guyon, Adrien Pavao, Kristin P Bennett, et al. Privacy preserving synthetic health data. In ESANN 2019-European Symposium on Artificial Neural Networks, Computational Intelligence and Machine Learning, 2019.

Daniel N Harris, Wei Song, Amol C Shetty, Kelly S Levano, Omar Cáceres, Carlos Padilla, Víctor Borda, David Tarazona, Omar Trujillo, Cesar Sanchez, et al. Evolutionary genomic dynamics of peruvians before, during, and after the inca empire. Proceedings of the National Academy of Sciences, 115(28):E6526–E6535, 2018.

Elena V Feofanova, Han Chen, Yulin Dai, Peilin Jia, Megan L Grove, Alanna C Morrison, Qibin Qi, Martha Daviglus, Jianwen Cai, Kari E North, et al. A genome-wide association study discovers 46 loci of the human metabolome in the hispanic community health study/study of latinos. The American Journal of Human Genetics, 107 (5):849–863, 2020.

Helgi Hilmarsson, Arvind S Kumar, Richa Rastogi, Carlos D Bustamante, Daniel Mas Montserrat, and Alexander G Ioannidis. High resolution ancestry deconvolution for next generation genomic data. bioRxiv, pages 2021–09, 2021.

Anders Boesen Lindbo Larsen, Søren Kaae Sønderby, Hugo Larochelle, and Ole Winther. Autoencoding beyond pixels using a learned similarity metric. In International conference on machine learning, pages 1558–1566. PMLR, 2016.

Diederik P Kingma and Max Welling. Auto-encoding variational bayes. arXiv preprint 1312.6114, 2013.

Scott Kirkpatrick, C Daniel Gelatt Jr, and Mario P Vecchi. Optimization by simulated annealing. science, 220(4598):671–680, 1983.

